# Vaccinia virus subverts xenophagy through phosphorylation and nuclear targeting of p62

**DOI:** 10.1101/2021.04.15.439938

**Authors:** Melanie Krause, Artur Yakimovich, Janos Kriston-Vizi, Moona Huttunen, Jason Mercer

## Abstract

Autophagy is an essential degradation program required to maintain cell homeostasis. Amongst its many functions is the engulfment and destruction of cytosolic pathogens, termed Xenophagy. Not surprisingly, many pathogens have adapted various strategies to circumvent or co-opt autophagic degradation during infection. For poxviruses, it is known that infection activates autophagy, which however is not required for successful replication. Despite the fact that these complex viruses replicate exclusively in the cytoplasm, autophagy-mediated control of poxvirus infection has not been explored. Using the prototypic poxvirus, vaccinia virus (VACV), we show that over-expression of the xenophagy receptors p62, NDP52 and Tax1Bp1 restricts poxvirus infection. While NDP52 and Tax1Bp1 were degraded, p62 was found to initially target cytoplasmic virions before being shunted to the nucleus. Nuclear translocation of p62 during infection was dependent upon p62 NLS2, and VACV kinase mediated phosphorylation of associated p62 residues T269/S272. These results indicate that VACV actively targets the xenophagy receptor p62 during the early stages of infection to avoid destruction, and further suggest that poxviruses exhibit a unique multi-layered control of autophagy in order to facilitate cytoplasmic replication.

## INTRODUCTION

Cell autonomous immunity represents the first line of defence used by cells to combat incoming pathogens (Randow, MacMicking, & James, 2013). Amongst the various strategies employed for detection and elimination of microbial invaders is xenophagy(Levine, 2005; Nakagawa et al., 2004a; Ohsumi, 2014). This selective form of macroautophagy(Feng, He, Yao, & Klionsky, 2014; Ravikumar et al., 2009), acts as a cytosolic defence mechanism for the detection, targeting, engulfment and delivery of cytoplasmic pathogens to lysosomes for degradation(Dong & Levine, 2013; Orvedahl & Levine, 2008). For the most part xenophagy uses the core autophagy machinery for the destruction of multiple pathogens, and relies on a sub-class of autophagy receptors for pathogen detection, namely the sequestome 1 (p62)-like receptors (SLRs): p62, NBR1, NDP52, Tax1Bp1 and optineurin (OPTN), (Dong & Levine, 2013; Sagar B. & Levine, 2009).

First described for bacteria(Gutierrez et al., 2004; Nakagawa et al., 2004b; Ogawa et al., 2005), xenophagy is also known to play an antiviral role through targeted degradation of cytosolic viruses or viral components (virophagy) or via activation of other cell autonomous antiviral responses(Dong & Levine, 2013). A wide range of viruses have been reported to be targeted by xenophagy including RNA viruses such as Influenza A, and DNA viruses including HSV-1 and HSV-2(Dong & Levine, 2013; Lee et al., 2010; Sun et al., 2012).

For vaccinia virus (VACV), a poxvirus which replicates exclusively in the cytoplasm of its host cells(Condit, Moussatche, & Traktman, 2006; Bernard Moss, 2007), evasion or disruption of xenophagy would seem to be of paramount importance. Initial studies aimed at investigating sequestration of autophagic membranes for VACV assembly showed that the absence of core autophagy components, ATG5 or beclin1, had no impact on VACV replication or infectious yield (Zhang et al., 2006). A subsequent investigation into the relationship between autophagy- and VACV-membrane biogenesis reported that VACV infection results in upregulation of LC3 lipidation independent of ATG5 and ATG7(Moloughney et al., 2011). The authors further demonstrate that VACV mediates aberrant ATG12-ATG3 conjugation and that late VACV infected cells are devoid of autophagosomes. As neither ATG3 nor LC3 lipidation were found to be required for productive infection, it was suggested that VACV might disrupt autophagy through aberrant LC3 lipidation and ATG12-ATG3 conjugation(Moloughney et al., 2011). Finally, a siRNA-based screen aimed at investigating the relevance of ATG proteins in viral replication found that VACV infection was enhanced upon depletion of SLRs important for xenophagy including p62, NDP52, NBR1 and OPTN (Mauthe et al., 2016). Taken together these studies suggest that VACV infection activates autophagy and that VACV either disarms or circumvents this host defense at multiple levels.

Here we sought to gain a better understanding of how VACV circumvents xenophagy by investigating its interplay with SLRs. Overexpression of the known xenophagy receptors revealed that p62, NDP52 and Tax1Bp1 could exert some control over VACV productive infection. While VACV appears to counter the effects of NDP52 and Tax1Bp1 through targeted degradation, we found that p62 was instead relegated to the nucleus of infected cells. Molecular dissection of p62 indicated that its NLS2 was necessary and sufficient for VACV-mediated nuclear shunting of p62. Furthermore, we show that the two VACV-encoded kinases, as opposed to cellular kinases, contribute to phosphorylation of p62 residues that serve to increase p62 NLS2 nuclear import activity. These results show that VACV actively avoids targeting by multiple SLRs during infection, uncover a new immunomodulatory role of virus-encoded kinases and suggest that poxviruses control xenophagy at several levels to assure successful cytoplasmic replication.

## RESULTS

### VACV circumvents autophagic restriction through NDP52, p62 and Tax1Bp1 targeting

While the role of autophagy in VACV infection remains largely undefined, a directed siRNA screen of ATG proteins indicated that inhibition of autophagy was beneficial to VACV replication(Mauthe et al., 2016). The SLRs: p62, NDP52, NBR1 and optineurin (OPTN) were amongst the strongest hits in this screen suggesting to us that this subset of autophagy receptors may serve in the autophagic elimination of VACV. While protein depletion studies are useful for defining cellular factors required for VACV infection(Beard et al., 2014; Kilcher et al., 2014; Moser, Jones, Thompson, Coyne, & Cherry, 2010; Sivan et al., 2013), defining VACV restriction factors is often more difficult due to the viruses capacity to perturb host cell innate immune responses(Bahar, Graham, Stuart, & Grimes, 2011; Bidgood & Mercer, 2015; Sivan et al., 2013). Thus to investigate the capacity of SLRs to restrict VACV replication we overexpressed GFP-tagged versions of the five known SLRs: NBR1, NDP52, p62, OPTN and Tax1Bp1 (Lazarou et al., 2015).

Transfected cells were infected with VACV and the infectious yield determined at 24 hours post infection (hpi). Overexpression of NDP52, p62 and Tax1Bp1 restricted VACV production by 92%, 82% and 75 % respectively, while overexpression of NBR1 and OPTN had no effect (Fig 1A). Immunoblot analysis confirmed that all SLRs were overexpressed to at least 50% of endogenous protein levels (Fig 1B). These results show that NDP52, p62 and Tax1Bp1 have the ability to restrict VACV infection when overexpressed, suggesting to us that VACV must have an intrinsic ability to overcome SLR-mediated restriction at native expression levels.

**Figure 1:**
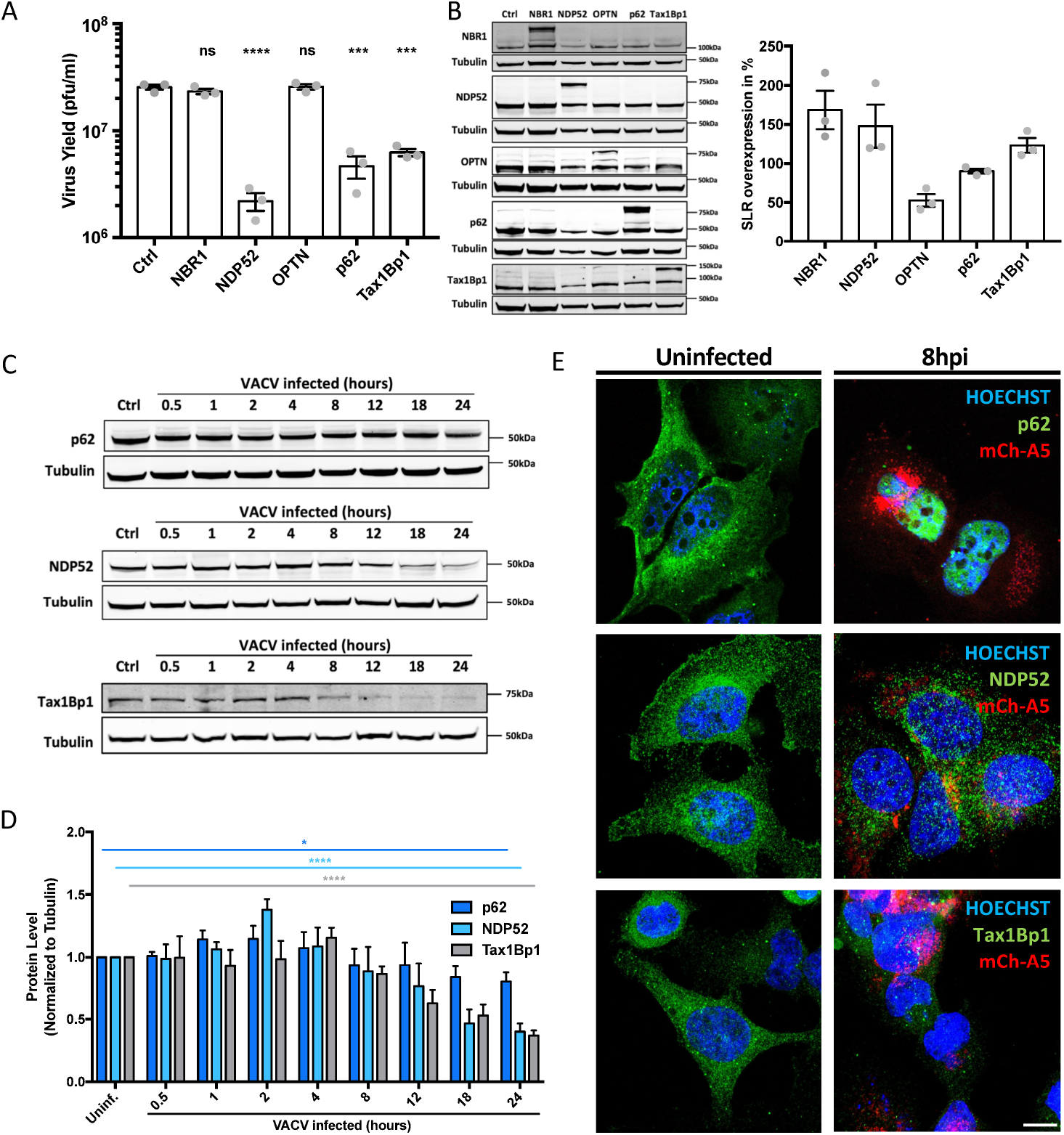
VACV interferes with p62-, NDP52- and Tax1Bp1-mediated viral restriction. A) 24h yield of WT VACV from HeLa cells overexpressing the indicated SLRs. Yields were determined by plaque assay on BSC40 cells. B) Quantification of SLR overexpression at 18 h post transfection. Representative immunoblot (left) and quantification (right). C) Assessment of p62, NDP52 and Tax1Bp1 protein levels during a time course of VACV infection. Representative oblots. D) Quantification of C. E) Assessment of p62, NDP52 or Tax1Bp1 localization by IF in uninfected and VACV mCh-A5 infected HeLa cells at 8 hpi. SLR (green), VACV mCh-A5 (red), Nuclei (blue). Representative images. All experiments were performed in triplicate and results displayed as mean ± SEM. Unpaired T-test with *P<0.05, ***P≤0.001 and ****P≤0.0001. Scale bars = 10 μM.

When we monitored the expression of these SLRs during the course of infection we found that both NDP52 and Tax1Bp1 levels were reduced over time, whereas p62 proteins levels remained largely unchanged during the course of infection (Fig 1C,D). Quantification indicated that the levels of both NDP52 and Tax1Bp1 were reduced from 8 hpi, culminating in less than 50% and 60% respectively by 24 hpi, and that p62 protein levels was reduced only slightly by <20% (Fig 1D). An immunofluorescence time course of cells infected with a VACV recombinant, VACV mCH-A5, that packages a mCherry-tagged version of the core proteins A5(Schmidt, Bleck, Helenius, & Mercer, 2011), and stained for NDP52, p62 or Tax1Bp1 confirmed the decrease in NDP52 and Tax1Bp1 protein levels compared to controls (Fig 1E and S1). Although we found only a minor decrease in p62 protein levels, strikingly p62 - which is usually dispersed throughout the cell - had been largely re-localized to the nucleus during VACV infection (Fig 1E). These results suggest that VACV overcomes SLR-mediated restriction by affecting cellular NDP52 and Tax1Bp1 protein levels, possibly through cleavage or degradation, and by shunting p62 to the nucleus.

### Nuclear targeting of p62 requires VACV early gene expression

VACV has been reported to direct proteasome-mediated disposal of cellular proteins including NDP52 and Tax1Bp1, while p62 was largely unaffected (Soday et al., 2019). Having shown that VACV core proteins are modified by K48-linked ubiquitin during assembly to facilitate core uncoating (Mercer et al., 2012), we reasoned that VACV drives p62 into the nucleus to avoid ubiquitin-mediated SLR recognition and subsequent autophagic degradation. To determine if incoming VACV cores could be recognized by p62, cells were infected with VACV mCh-A5 and subsequently stained for p62 at 2 hpi (Fig 2A). While we found instances of p62 ring-formation around cytoplasmic VACV cores, this was a rare event suggesting that VACV can avoid p62-targeting early in infection.

**Figure 2:**
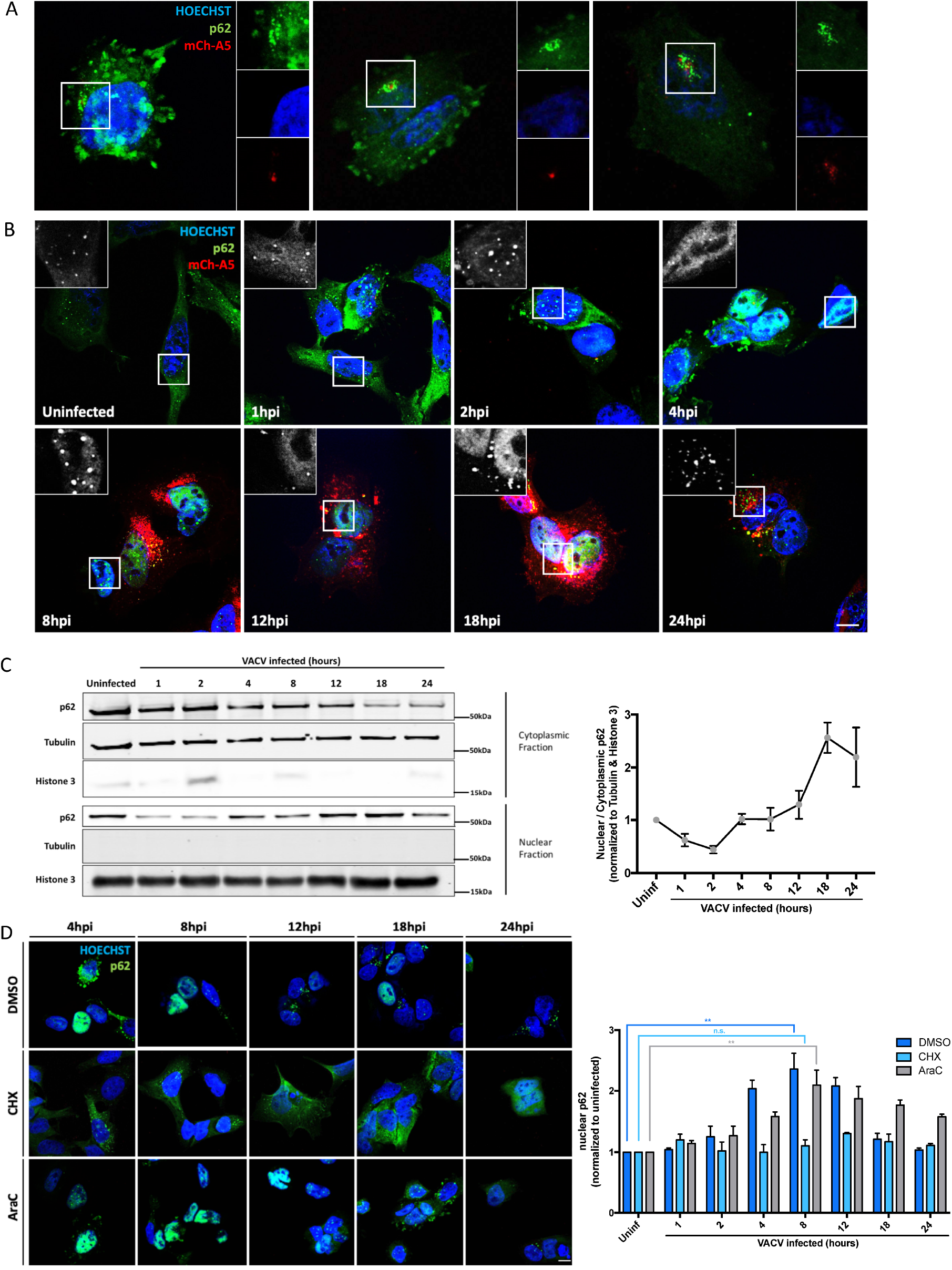
VACV evades p62 via early gene-mediated nuclear relocation. A) Incoming VACV virions are sometimes targeted by p62. HeLa cells infected with VACV mCh-A5 (red), were fixed and immunostained for p62 (green) and nuclei (blue) at 2 hpi. B) HeLa cells infected with VACV mCh-A5 (red) were fixed and immunostained for p62 (green) and DNA (blue) at the indicated time points. C) HeLa cells, uninfected or infected WT VACV, were harvested at the indicated timepoints and subjected to nuclear/cytoplasmic fractionation followed by immunoblot analysis for p62. Tubulin and histone 3 served as quality controls for cytoplasmic and nuclear fractionation, respectively. Quantification of nuclear/cytoplasmic p62 displayed to the right. D) HeLa cells infected with VACV WR mCh-A5 in the presence of DMSO, Cycloheximide (CHX) or Cytosine Arabinoside (AraC), were fixed and stained for p62 (green) and DNA (blue) at the indicated timepoints. Quantification of nuclear p62 displayed at right. All experiments were performed in triplicate and representative images displayed. Quantifications are presented as mean ± SEM in (C) or + SD (D). Unpaired T-test with **P≤0.01 and ns = non-significant. Scale bars = 10 μM.

To determine when p62 was translocated to the nucleus we carried out an image-based time course of infection (Fig 2B). In uninfected cells, p62 appeared to be distributed predominantly in the cytoplasm with some in the nucleus. The cytoplasmic signal seemed to increase at 1 hpi, before p62 appeared to aggregate in the cytoplasm by 2 hpi. By 4 hpi p62 could be predominantly detected in the nucleus of infected cells where it then resided over the course of infection until 24 hpi, when only low levels of p62 could be detected as small aggregates in the cytoplasm (Fig 2B). We confirmed these results using nuclear/cytoplasmic fractionation of infected cells combined with immunoblots for p62, as well as tubulin and Histone 3 which served as fractionation controls. Consistent with the imaging results p62 was found mostly in the cytoplasm, but also in the nucleus of uninfected cells (Fig 2C). Using this as a baseline, we quantified the nuclear/cytoplasmic distribution of p62 during VACV infection (Fig 2C; right panel). By 1 hpi the distribution of p62 shifted to cytoplasmic where it remained until 4 hpi. At this time, p62 distribution shifted back to the nucleus where it accumulated until 18 hpi, before equalizing between the two compartments (Fig 2C).

That VACV infection initiated p62 aggregation and nuclear translocation by 2 hpi suggested that a VACV early protein(s) was responsible for this effect. To test this, HeLa cells infected with VACV in the presence of the translation inhibitor cycloheximide (CHX) or Cytosine arabinoside (AraC), which blocks VACV DNA synthesis, were assessed for nuclear p62 (Fig 2D). As expected in untreated, infected cells p62 was found predominantly in the nucleus between 4-24 hpi. In the presence of CHX, which blocks early viral protein synthesis (B. Moss & Filler, 1970), p62 did not accumulate in nuclei. In the presence of AraC, nuclear accumulation of p62 was delayed (Fig 2D; right panel). As AraC prevents intermediate and late gene expression, without impacting early genes (Furth & Cohen, 1968), these results suggest that VACV early gene(s) initiates p62 nuclear translocation and that one or multiple late genes facilitates maintenance of this phenotype.

### Nuclear localization signal 2 of p62 is required for VACV-mediated nuclear shuttling

We next asked how VACV mediates nuclear translocation of p62. Of relevance, p62 contains two nuclear localization signals (NLSs); NLS1 (aa 186-189) and NLS2 (aa 264-267). Mutation of the two basic residues in NLS1 or NLS2 (illustrated in Fig 3A) showed a 1.5- and 6.3-fold defect in nuclear import, respectively, suggesting the NLS2 is the predominant NLS required for p62 import (Pankiv et al., 2010). To test if either NLS was important for VACV-mediated nuclear translocation, EGFP-tagged versions of p62, p62 NLS1^mut^ or p62 NLS2^mut^ were transfected into HeLa cells. Immunoblot analysis assured equivalent expression levels of the three constructs (Fig S2). Transfected cells were either left uninfected or were infected with VACV mCh-A5 and assessed for p62 nuclear translocation at 4 and 8 hpi. As expected, in uninfected cells WT p62 appeared to be evenly distributed while both NLS1^mut^ and NLS2^mut^ p62 showed greater cytoplasmic accumulation (Fig 3B). By 4 hpi both WT p62 and p62 NLS1^mut^ had largely re-localized to nuclei, while the localization of p62 NLS2^mut^ remained unchanged. A similar localization pattern was seen at 8 hpi. Quantification of the relative intensity of the nuclear EGFP signal in VACV infected cells showed a 5-fold increase in WT p62 nuclear signal by 8 hpi (Fig 3C). Conversely, the p62 NLS2^mut^ showed no increase in nuclear localization upon VACV infection at either 4 or 8 hpi. The p62 NLS1^mut^ displayed an intermediate phenotype, its nuclear signal increased 3.5-fold by 8 hpi (Fig 3C). These results indicated that NLS2 is required for VACV-mediated nuclear re-localization of p62 during infection. Consistent with previous reports, mutation of NLS1 did not prevent, but impaired the efficiency of p62 nuclear shuttling.

**Figure 3:**
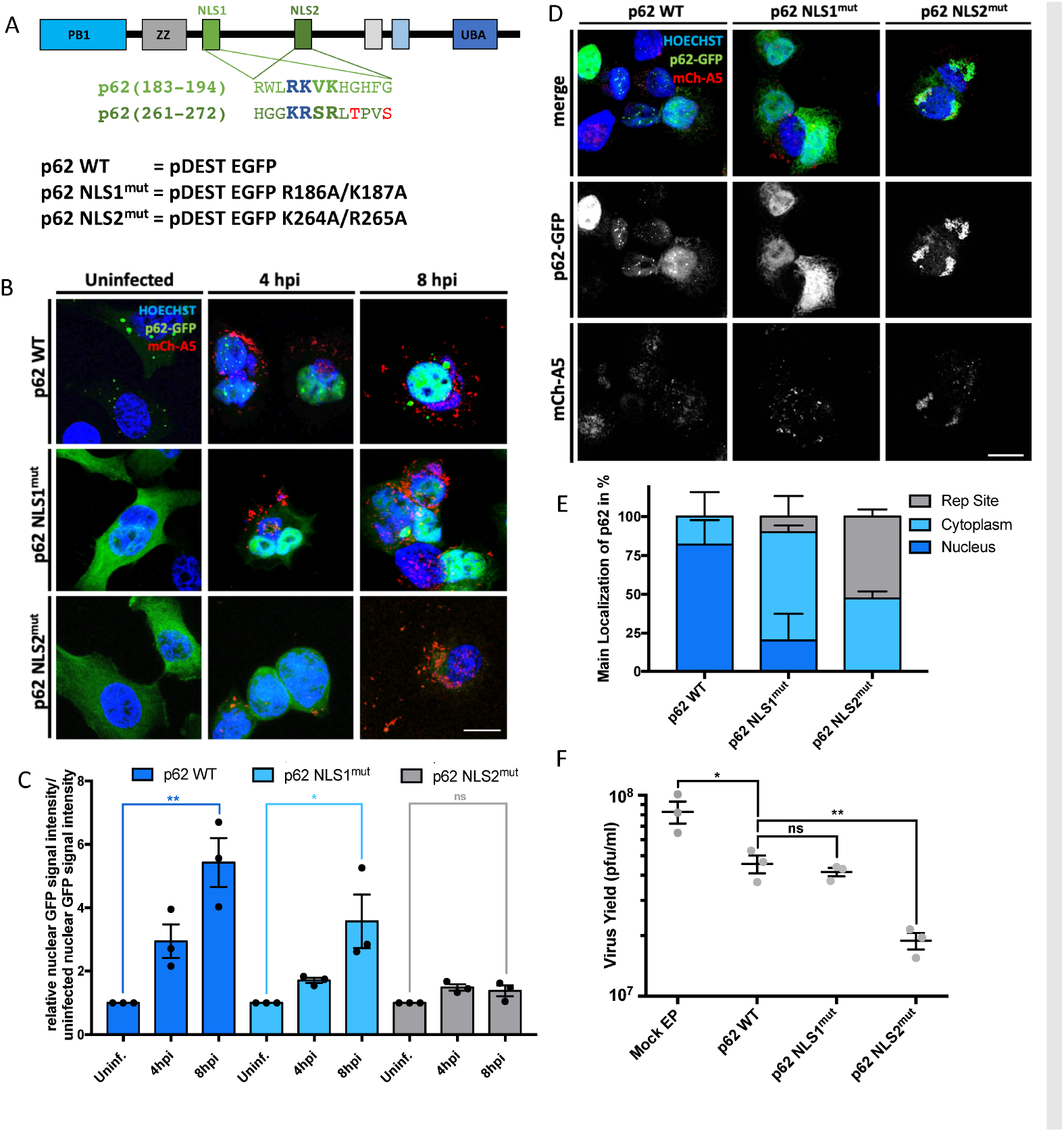
VACV-mediated nuclear shunting of p62 depends on p62 NLS2. A) Top: Schematic of the two p62 NLS sites. Functional NLS motifs bolded and enlarged. Blue letters indicate amino acids changed in NLS1^mut^ and NLS2^mut^. Bottom: EGFP-tagged p62 constructs WT, NLS1^mut^ and NLS2^mut^ with indicated mutations B) HeLa cells expressing the indicated EGFP-p62 proteins (green) were infected with VACV mCh-A5 (red) and fixed at 4 or 8 hpi. C) Quantification of nuclear WT, NLS1^mut^ and NLS2^mut^ p62 signals from B. D) HeLa cells expressing WT, NLS1^mut^ or NLS2^mut^ p62 were infected with VACV mCh-A5 (red), fixed at 8 hpi and cells immunostained with anti-GFP (green) E) Quantitative classification of p62 localization from images in D. E) 24 h yield of WT VACV from cells expressing EGFP tagged WT, NLS1^mut^ or NLS2^mut^ p62. All experiments were performed in triplicate (n ≥ 50 cells/construct) and representative images displayed. Quantifications are presented as mean ± SEM. Unpaired T-test with *P<0.05 and **P≤0.01. A) is adapted and modified from Pankiv *et al*., 2009.

### Cytoplasmic p62-NLS2^mut^ reduces virus yield and localizes to VACV replication sites

Given the impact of p62 overexpression on viral yield (Fig 1A,B), we wondered if the cytoplasmic retention of p62 NLS2^mut^ would impact VACV production. For this, cells transfected with plasmids expressing EGFP, WT p62, p62 NLS1^mut^ or p62 NLS2^mut^ were infected with WT VACV. At 24 hpi cells the productive infectious virus yield was determined by plaque assay (Fig 3F). As before, over expression of WT p62 reduced virus production by 45%. While overexpression of p62 NLS1^mut^ impacted virus yield to a similar level (50%), overexpression of p62 NLS2^mut^ reduced virus yield an additional 27% over WT and NLS1^mut^ p62 overexpression for a total reduction of 77% when compared to EGFP-expressing control cells (Fig 3F).

That overexpression of the p62 NLS2^mut^ was detrimental to productive VACV infection suggested that cytoplasmic p62 might directly target assembling VACV virions. As VACV core proteins are ubiquitinated during assembly, we reasoned that cytoplasmic p62 would target these proteins within VACV replication sites. In order to accurately assign the localization of EGFP WT p62 and p62 NLS2^mut^ in VACV infected cells, anti-EGFP antibody was used to amplify the cytoplasmic EGFP-p62 signal (Fig 3D). As expected, WT p62 localized to nuclei in 82% of infected cells at 8 hpi and to the cytoplasm in the other 18% (Fig 3E). In the case of NLS2^mut^, p62 was found in the cytoplasm in all cells. Strikingly, in 53% of these cells, p62 was predominantly associated with VACV replication sites (Fig 3D). Collective these results strongly suggest that VACV shunts p62 to the nucleus to prevent autophagic degradation of nascent virions.

### VACV infection upregulates p62 (T269/ S272) phosphorylation independent of p38

Multiple studies have identified T269/S272 as two phosphorylation sites near p62 NLS2 (Nousiainen, Silljé, Sauer, Nigg, & Körner, 2006; Olsen et al., 2006; Pankiv et al., 2010; Yanagawa, Yuki, Yoshida, Bannai, & Ishii, 1997). Follow up work demonstrated that a phosphomimetic p62 T269E/S272E mutant was completely nuclear, leading the authors to conclude that T269/S272 phosphorylation serves to regulate p62 NLS2-dependent nuclear localization (Pankiv et al., 2010). Considering the robustness of VACV-mediated p62 nuclear shunting, we asked if VACV infection increases p62 T269/S272 phosphorylation. Cells were infected with WT VACV and p62 T269/S272 phosphorylation assessed at various timepoints using a p62 T269/S272 phospho-specific antibody (Fig 4A; top). Increased p62 T269/S272 phosphorylation was seen by 1 hpi and remained up-regulated until 18 hpi before returning to background levels (Fig 4A; bottom).

**Figure 4:**
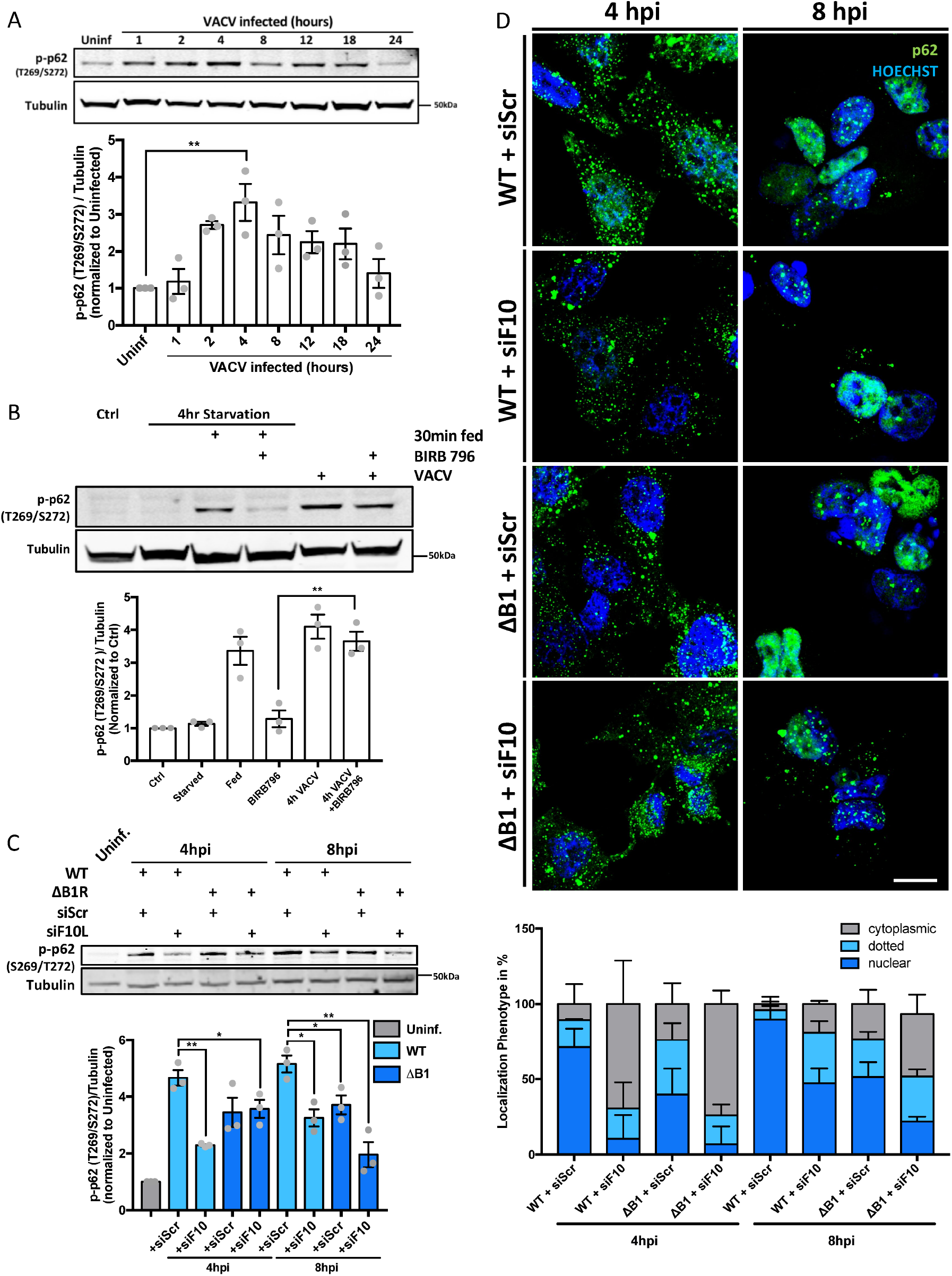
VACV encoded kinases B1 and F10 phospho-regulate p62 nuclear shunting. A) Top: Immunoblot analysis of p62 (Thr269/Ser272) phosphorylation during VACV infection. Bottom: Quantification of p-p62 relative to uninfected controls. B) Top: A549 cells, untreated or treated with the p38δ inhibitor BIRB796 (10 µM), were infected with VACV and p-p62(Thr269/Ser272) assessed at 4 hpi. Cells starved for 4 h then fed in the absence or presence of BIRB796 served as a positive control for inhibition of p38δ-mediated p62 phosphorylation. Bottom: Quantification of p-p62(Thr269/Ser272). C) Top: Immunoblot analysis of HeLa cell lysates infected with WR WT or WR ΔB1 virus in the absence or presence of VACV F10-targeting siRNA. Top: Cells were harvested at 4 and 8 hpi and subjected to immunoblot analysis for p-p62(Thr269/Ser272). Bottom: Quantification of p-p62(Thr269/Ser272). D) HeLa cells transfected with scrambled or F10-targeting siRNA were infected with WT or ΔB1 WR VACV, fixed at 4 or 8 hpi and stained for p62 (green) and DNA (blue). Bottom: Quantification of p62 localization phenotype (cytoplasmic, dotted or nuclear) at 4 and 8 hpi. All experiments were performed in triplicate and representative images displayed. Quantifications are presented as mean ± SEM. Unpaired T-test with *P<0.05 and **P≤0.01.

Of relevance, p38δ (mitogen activated protein kinase 13)-mediated phosphorylation of p62 T269/S272 has been shown to regulate mTORC1-mediated inhibition of autophagy (Linares et al., 2015). To determine if p62 T269/S272 phosphorylation during VACV infection was via p38δ, cells were either starved or infected in the absence or presence of BIRB796, an allosteric pan-p38 MAPK inhibitor (Escós, Risco, Alsina-Beauchamp, & Cuenda, 2016). When p62 T269/S272 phosphorylation assessed by immunoblot, BIRB796 was found to effectively prevented starvation-induced p62 phosphorylation while no significant difference in VACV-mediated p62 phosphorylation was observed in the absence or presence of BIRB796 (Fig 4B). These results indicate that VACV does not act through p38δ to phosphorylate p62 and suggested to us that a viral kinase may be responsible for p62 phosphorylation.

### VACV kinases are required for p62 phosphorylation and nuclear translocation

VACV encodes two kinases, B1 and F10, both of which are essential for viral propagation (Lin & Broyles, 1994; Traktman, Caligiuri, Jesty, Liu, & Sankar, 1995; Wang & Shuman, 1995). B1 is a S/T kinase expressed early during infection and required for viral DNA replication(Jamin, Ibrahim, Wicklund, Weskamp, & Wiebe, 2015). F10 is a late expressed S/T/Y kinase required for virus assembly (Szajner, Weisberg, & Moss, 2004). While no deletion virus is available for F10, a B1 deletion virus (ΔB1R) has been recently generated (Olson, Rico, Wang, Delhon, & Wiebe, 2017).

Using the ΔB1R virus and a siRNA directed against F10, we asked if either B1 or F10 contribute to p62 phosphorylation at 8 hpi, a time point when the F10 siRNA efficiently reduced expression of F10 protein (Fig S3). Cells transfected with control or F10 siRNA were infected with WT or ΔB1R VACV and p62 S269/T272 phosphorylation determined (Fig 4C). At 8hpi WT VACV infection increased p62 S269/T272 phosphorylation by 5.1-fold over uninfected cells. This was reduced to 3.2- and 3.7-fold in the absence of either F10 or B1 respectively and to 1.9-fold when both kinases are absent.

As these results suggested that both B1 and F10 contribute p62 phosphorylation we next asked how loss of these kinases would affect p62 nuclear translocation during VACV infection. Cells transfected with control or F10 siRNA were infected with WT or ΔB1R VACV and the cellular distribution of p62 monitored by immunofluorescence at 4- and 8 hpi (Fig 4D, top). As expected, in the vast majority (90%) of control siRNA transfected cells infected with WT VACV, p62 was found within the nucleus (Fig 4D; bottom). Depletion of F10 and deletion of B1 led to a 37% and 41% reduction in p62 nuclear translocation, respectively. In the absence of both B1 and F10, the number of cells displaying VACV-mediated p62 nuclear translocation was reduced to 21.8% (Fig 4D, bottom). Collectively, these results support a role for both VACV encoded kinases, B1 and F10, in phosphorylation of p62 and its subsequent nuclear translocation.

## DISCUSSION

As a virus that replicates exclusively in the cytoplasm of host cells VACV is subject to a battery of cell autonomous immune defences (Bidgood & Mercer, 2015; Hu & Shu, 2018, 2020). To overcome this VACV dedicates nearly half of its 200 encoded proteins to evading host cell defences and subjugating host cell systems (Bahar et al., 2011; Bidgood & Mercer, 2015; Lu & Zhang, 2020). Cellular degradation pathways are no exception. Manipulation of the ubiquitin proteasome system, for example, has been shown to be essential for VACV genome uncoating and targeted degradation of viral restriction factors (Mercer et al., 2012; Soday et al., 2019). We previously demonstrated that during cytoplasmic virus assembly VACV core proteins are ubiquitinated and packaged into nascent virions. Subsequently, during virus entry the ubiquitinated cores are released into the cytoplasm where they direct proteasome-mediated core degradation and genome release (Mercer et al., 2012).

There have also been several investigations into the role of autophagy in VACV membrane biogenesis: it was shown that VACV activates autophagy and induces aberrant ATG12-ATG3 conjugation, that VACV infected cells are devoid of autophagosomes late in infection, that VACV replication does not require LC3 lipidation, ATG3 or ATG5. In addition, it was reported - in a large-scale screen - that RNAi-mediated depletion of SLRs enhances VACV infection (Mauthe et al., 2016). Given that all SLRs use ubiquitin for cytosolic pathogen recognition and destruction (Deretic, Saitoh, & Akira, 2013; Dupont et al., 2009; Thurston, 2009; Wild et al., 2011), and we know that ubiquitinated viral cores are exposed to the cytoplasm during two stages of the VACV lifecycle - entry and assembly - we set out to investigate the interplay between VACV, SLRs and xenophagy.

Overexpression studies revealed that p62, NDP52 and Tax1Bp1 have the ability to restrict VACV productive infection by 90%, 62% and 58%, respectively. This indicated that xenophagy can restrict VACV infection, but under normal conditions VACV is able to overcome this. When we followed these proteins over a time-course of infection we found that both NDP52 and Tax1Bp1 protein levels were reduced. This suggested to us that VACV either directs the cleavage or degradation of these proteins. VACV encodes two proteases, G1 and I7 (Ansarah-Sobrinho & Moss, 2004b, 2004a). While G1 substrates and cleave specificity is unknown, I7 cleaves specifically at AGX sites in several viral structural proteins and has been shown to cleave the antiviral protein Dicer (Byrd, Bolken, & Hruby, 2002; Chen et al., 2015; Novy et al., 2018). However, no AGX sites are present in either NDP52 or Tax1Bp1, and inhibition of I7 and G1 expression had no impact on reduction of NDP52 or Tax1Bp1 protein levels during infection (data not shown). This suggested to us that VACV mediates the degradation of these two SLRs. Consistent with this, proteomic analysis of VACV infected cells, suggests that both NDP52 and TaxBp1 are subject to proteasome-mediated degradation(Soday et al., 2019).

As opposed to NDP52 and TaxBp1 we found that p62 remained relatively stable over the course of infection suggesting that VACV employed an alternative strategy to disable or evade this SLR. Consistent with their being ubiquitinated, we found that a small subset (<5%) of incoming virions could be targeted by cytoplasmic p62. We noted that upon infection the distribution and subsequently the cellular localization of p62 changed from diffuse and cytoplasmic to aggregated and ultimately nuclear. Although generally considered a cytoplasmic protein, p62 is known to shuttle between the cytoplasm and the nucleus. Pankiv *et al*. first reported that p62 contains two NLS motifs (NLS 1 and NLS2) of which NLS2 appears to be the more important one (Pankiv et al., 2010). Our molecular dissection of p62 showed that NLS2 was indeed largely responsible for p62 nuclear shuttling. In the absence of NLS2, p62 was found at cytoplasmic VACV replication sites, presumably due to targeting of ubiquitinated core proteins, and virus production was reduced.

In addition to autophagy p62 is involved in regulating cellular processes such as oxidative stress responses, NFκB signalling and mTORC1 activity (Ichimura et al., 2013; Jain et al., 2010; Katsuragi, Ichimura, & Komatsu, 2015; Linares et al., 2015; Martin, Diaz-Meco, & Moscat, 2006; Moscat, Diaz-Meco, Albert, & Campuzano, 2006; Sanchez-Garrido, Sancho-Shimizu, & Shenoy, 2018). Not surprisingly each of these pathways intersect with and are regulated by VACV (Meade et al., 2018; Smith et al., 2013). Of relevance to our study mTORC1 inhibits autophagy when it is ubiquitinated by TRAF6 (Linares et al., 2013). This E3 ubiquitin ligase is recruited to mTORC1 by p62 after phosphorylation of p62 residues T269/S272 by p38δ (Linares et al., 2015). Pankiv *et al*. saw that the nuclear import activity of NLS2 was modulated by phosphorylation of these same residues (Pankiv et al., 2010). We found that VACV infection resulted in increased p62 T269/S272 phosphorylation, raising the possibility that VACV might be exerting its control over p62 localization through p38δ. However, using a p38δ inhibitor we found no evidence of p38δ involvement in p62 phosphorylation. This finding was consistent with reports that VACV-mediated dysregulation of mTOR protein biosynthesis activity is uncoupled from its control over autophagic responses (Meade, King, Munger, & Walsh, 2019).

In the absence of p38δ involvement, we turned to the two VACV-encoded kinases B1 and F10 (Kovacs, Vasilakis, & Moss, 2001; Wang & Shuman, 1995). B1 and F10 are expressed temporally; the former being an early gene is expressed upon virion entry prior to virion uncoating while F10 being is a late gene, expressed after DNA replication. That p62 nuclear translocation was largely prevented in the absence of early gene expression and delayed in the absence of late gene expression, suggested to us that both B1 and F10 may play a role in this process. Deletion of B1 and depletion of F10 indicated that both kinases played a role in p62 phosphorylation and nuclear translocation. Given the temporal nature of their expression it reasons that B1 initially drives p62 to the nucleus to prevent targeting of incoming ubiquitinated cores, and F10 serves to maintain this phenotype during viral morphogenesis in order to hamper xenophagy of nascent ubiquitinated core proteins prior to packaging. These results suggest that in addition to B1-mediated regulation of BAF (Wiebe & Traktman, 2007), VACV encoded kinases may play a larger role in immune modulation that previously appreciated.

In sum, using over expression of SLRs we have shown that xenophagy can impart control over VACV productive infection. In support of this, infection with a highly attenuated strain of vaccinia - modified vaccinia Ankara (MVA) - has been reported to result in the induction of autophagy (Tappe et al., 2018). This phenomenon is largely masked during normal infection, which we attribute to VACVs ability to effectively disarm the xenophagy receptors NDP52, p62 and Tax1Bp1. Exemplifying the ability of poxvirus to exert a multi-layered control over cell intrinsic immune responses, we show that VACV uses distinct mechanisms - degradation of NDP52 and TaxBp1 versus cytoplasmic expulsion of p62 - to assure that none of these SLRs can contribute to xenophagy. Interestingly, it was recently shown that nuclear p62 plays a role in trafficking of NFκB and aggresome-related proteins to nucleolar aggresomes in order to suppress stress-induced apoptosis (Lobb et al., 2021). Given the extensive regulation of NFκB by VACV(Smith, Talbot-Cooper, & Lu, 2018), it will be of future interest to investigate possible pro-viral role(s) of nuclear p62 during infection.

## MATERIALS AND METHODS

### Cells and Viruses

BSC40 cells, A549 cells and HeLa cells were maintained in Dulbecco’s Modified Eagle Medium (DMEM, Life Technologies) supplemented with 10 % FBS, 1 mM sodium pyruvate, 100 μM non-essential amino acids, 2 mM L-alanyl-L-glutamine dipeptide and 1 % Penicillin-Streptomycin under at 37°C and 5 % CO2. Cell lines passaged two to three times per week using PBS and Trypsin/EDTA and tested frequently for mycoplasma. VACV WR WT, WR E EGFP(Chomczynski & Mackey, 1995; Stiefel et al., 2012), WR L EGFP(Chomczynski & Mackey, 1995; Schmidt et al., 2013; Stiefel et al., 2012), WR mCherry-A4(Mercer & Helenius, 2008; Schmidt et al., 2013) were previously published. WR ΔB1mutB12(Olson, Wang, Rico, & Wiebe, 2019; Rico et al., 2019) was a kind gift from the Wiebe lab.

### Reagents

Cytosine arabinoside (AraC) and cycloheximide (CHX) were obtained from Sigma-Aldrich. drugs were used at 10 μM and 50 μM respectively. DMSO (Sigma-Aldrich) was used to dissolve drugs and as negative control in all drug related assays. p38 MAP Kinase Inhibitor BIRB 796 (MERCK Millipore) was used at 10 μM. Anti-GFP(Kilcher et al., 2014) antibody was used at 1:500 in WB. The following antibodies were purchased from CST and used at concentrations indicated in brackets: Histone 3 (#9701S; WB 1:5000), NDP52 (#60732S; WB 1:1000; IF 1:100), p-p62 (Thr269/Ser272) (#13121S; WB 1:1000), Tax1Bp1 (#5105S; WB 1:1000; IF 1:100), Tubulin (#9701S; WB 1:5000). p62 antibody (#P0067; WB 1:5000; IF 1:1000) was purchased from Sigma. Hoechst Trihydrochloride Trihydrate 33342 (Invitrogen #H3570) was used for DNA staining at 1:10,000. IRDye-coupled secondary antibodies were purchased from Li-COR and used at 1:5000. Alexa Fluor-conjugated secondary antibodies and phalloidin were purchased from Invitrogen and used at 1:400. SLR-GFP constructs were kindly gifted by Richard Youle (NIH)(Lazarou et al., 2015). Mutant-p62 constructs were a kind gift from Terje Johansen (University of Tromsø)(Pankiv et al., 2010). F10L siRNA was custom designed and manufactured by Ambion Life Technologies (F10L: GAACUACCCUGUUGCGACAtt; F10L_as: UGUCGCAACAGGGUAGUUCgt).

### MV 24-hour yield

For 24 hr yields, 30 mm dishes of HeLa cells were infected in DMEM without FBS at MOI 1 or MOI 0.1 with WT VACV and incubated for 1 hr at 37°C. After incubation, infection media was aspirated and replaced with full DMEM, containing compounds of interest at concentrations previously indicated. Alternatively for yields of siRNA-transfected cells, cells were infected at 72 hrs post siRNA transfection, as described below, and incubated in full medium. After 24 hrs incubation or any other timepoint of interest, the media was aspirated and cells were scraped into 1 ml of PBS and spun at 300 x *g* for 5 min. The cell pellets were then resuspended in 100 μl 1 mM Tris pH 9.0 and freeze-thawed three times in liquid nitrogen. The obtained MV solution was used in a dilution series on confluent BSC40 cells for plaque assay analysis starting at 10^−4^ to 10^−9^.

### Plaque assay

To determine the titre of a purified virus stock or 24-hr yields, plaque assays were performed. Serial dilutions were carried out as indicated above. 500 μl of each dilution was applied to one well in a 6-well dish containing a monolayer of BSC-40s at 100 % confluency and 500 μl of DMEM without FBS. Cells were incubated for 1 hr at 37°C. Infection medium was then aspirated and replaced with full supplementary media containing FBS. After 48 hrs of incubation at 37°C, media was aspirated and the cells fixed with 0.1 % crystal violet and 2 % formaldehyde. Plaque-forming units per millilitre (pfu/ml) were calculated by manually counting of plaques. For experiments in other cell lines the MOI was adjusted. Since BSC40 cells are more permissive than HeLa cells, a ten times higher MOI is required in HeLa cells to match the infection level in BSC40 cells. An MOI of 30 for BSC40s is equivalent to an MOI of 3 in HeLa cells.

### Microscopy Assays

For confocal imaging HeLa cells were seeded on 13 mm glass coverslips (VWR) at 60,000 cells per coverslip the day prior to infection. Infection was carried out at MOI 10 in DMEM, and incubated for 1 hr at 37°C. The media was removed and replaced with full medium and incubated for the desired time. Cells were fixed with 4 % FA-PBS for 15 mins. Unless noted otherwise cells were permeabilised with ice cold MeOH at −20°C for 20 min, blocked with 3 % bovine serum albumin (BSA, from Sigma Aldrich) in PBS for 1 hr, and stained with 30 μl primary antibody in 3 % BSA-PBS for at least 1 hr at RT or overnight at 4°C. Secondary antibody staining was performed for 1 hr at RT. Samples were imaged using a 63 x oil immersion objective (ACS APO) on an Leica TCS 2012 model SPE confocal microscope. For high content imaging antibody and Hoechst staining was carried out in a 40 μl volume on a shaker as stated above. The Opera Phenix High Content Screening System was used for image acquisition at 40 x magnification with an air objective using 405 nm, 488 nm, 594 nm or 647 nm lasers and at least 15 images taken per well. CellProfiler was used to detect individual cells, based on Hoechst stained nuclei.

### Infection time courses

For western blot or fractionation samples, HeLa cells were seeded in either 60 mm or 35 mm dishes for confluency at infection. Infection was carried out in DMEM without FBS at MOI 30. VACV was incubated with the cells for 1 hr at 37°C, before aspirating and replacing with full medium. Cells were either left untreated or treated with the indicated compounds from the start of infection, and incubated at standard conditions. Samples were harvested at their respective timepoints, by removing media and washing cells with cold PBS and subsequently scraped into 50 – 200 μl (depending on dish size) lysis buffer containing protease inhibitor (#5872, NEB). The sample was then left on ice for at least 20 mins and subsequently spun down at 20,000 x g for 10 mins at 4°C. The supernatant was either frozen and stored at −20°C or directly supplemented with 3 x blue loading dye containing DTT. Prior to western blot analysis the samples were subject to 5 min incubation at 95°C. Protein samples were loaded into 12 % Bis-Tris polyacrylamide gels and transferred onto 0.2 μm nitrocellulose membrane. Membranes were blocked with 5 % BSA TBS-T. Primary antibodies were applied in 5 % BSA TBS-T over night at 4°C. Membranes were incubated with LiCor secondary antibodies for 1 hr at RT. Imaging of membranes was carried out with using a LiCor Odyssey. Western blot quantifications were done using ImageJ (Version2.0.0) with tubulin used as a loading control where applicable. For separation of nuclear and cytoplasmic fractions of cell lysates the Qproteome cell compartment kit (Invitrogen #37502) was used according to manufacturer protocol.

### DNA Electroporation

To express mutant cellular proteins in vitro, DNA electroporation was conducted following the Amanxa^®^ Cell line nucleofector protocol using the Cell Line Nucleofector^®^ Kit R (Lonza). 2 μg plasmid DNA was used per reaction and electroporation was carried out using the I-013 program for high expression efficiency of HeLa cells. Afterwards cells were immediately resuspended in 1.5 ml 37°C warm DMEM containing FBS and plated in a 6-well plate with one electroporation reaction per well. The cells were incubated over night at 37°C for ∼16 hrs before infection or harvesting for subsequent analysis.

### Cell Count Assay

For automated cell counting of p62 nuclear translocation a median filter was applied on the nuclear channel using a 3-pixel radius parameter followed by Otsu segmentation where the thresholding parameter was calculated from the image stack. A custom ImageJ macro calculated the number of nuclei using watershed segmentation algorithm was with noise tolerance parameter 110.

### Subcellular Intensity Measurement

Computational analysis of the image-based data from microscopy modalities has been performed using combination of opensource software and custom developed code. Image analysis was performed on a Desktop PC equipped with Intel Core i7-8700K CPU at 3.7 GHz and 32 GB of RAM as well as GeForce 1080 Ti GPU. To measure single cell intensities in confocal images stacks Z-maximum intensity projection was performed. Next, upon Gaussian smoothing cells were detected in a custom CellProfiler pipeline(Carpenter et al., 2006). Finally, single cell intensities were averaged per image and condition. ImageJ(Schindelin et al., 2012) was used to batch process the dataset.

## Acknowledgements

This research was supported by MRC LMCB PhD programme at University College London (MK), MRC LMCB Grant Ref. MC_UU_00012/7 (JM), the European Research Council 649101-UbiProPox (JM), and MRC core funding to the MRC-LMCB Grant Ref. MC_U12266B (JKV). We thank Richard J. Youle (National Institute of Neurological Disorders and Stroke, NIH) for the autophagy SLR plasmids and Terje Johansen (University of Tromsø) for the p62 NLS mutant plasmids.

## Supplementary Figures

**Figure S1:**
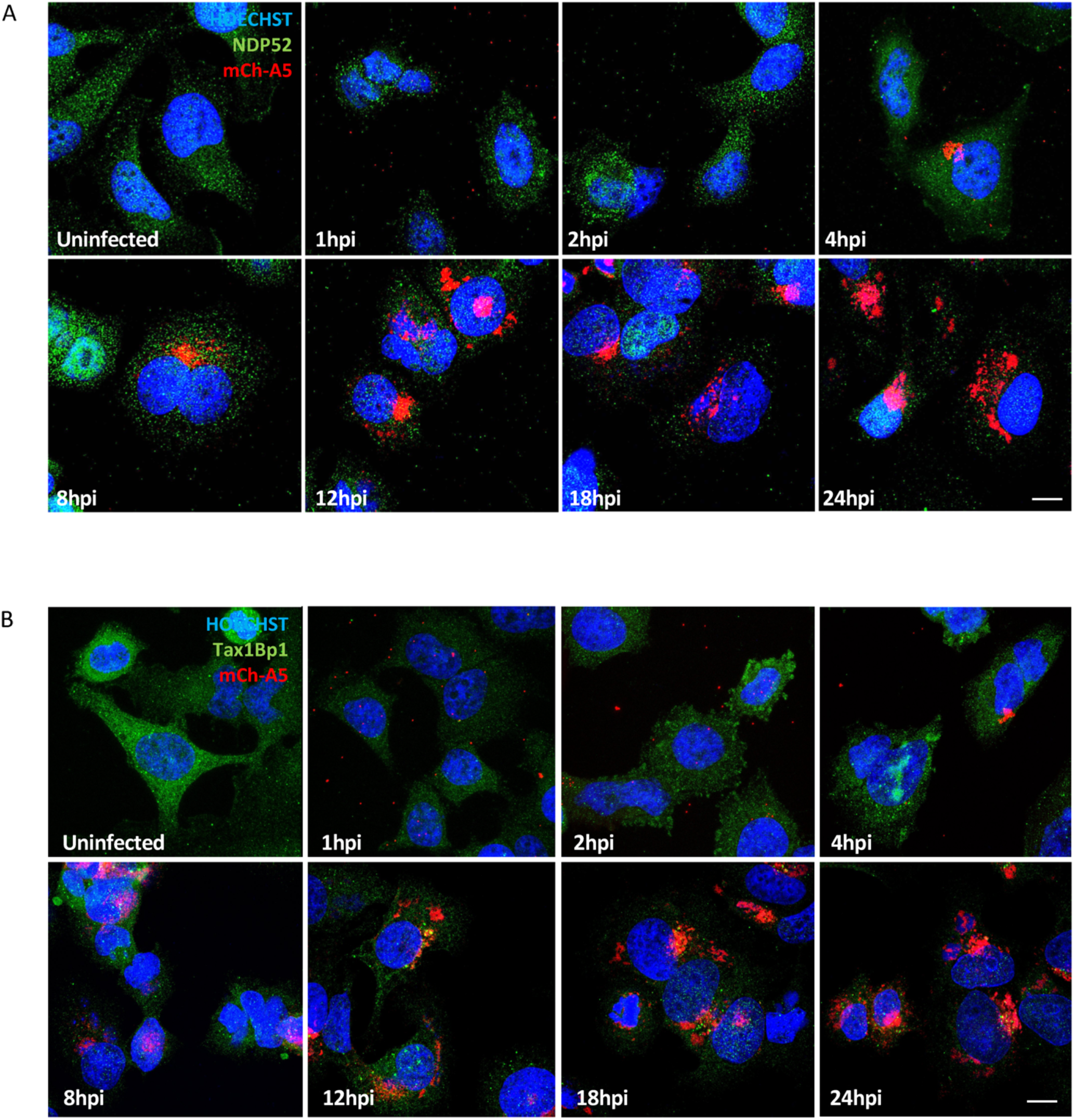
Abundance of autophagy receptors NDP52 and Tax1Bp1 seems reduced during VACV infection. A) Time course of HeLa cells infected with WR mCh-A5 (red) and immunostained for NDP52 (green) and stained for DNA (blue). B) Time course of HeLa cells infected with WR mCh-A5 (red) and immunostained for Tax1Bp1 (green) and stained for DNA (blue). Scale bars = 10 µM. Experiments performed in triplicate and representative images displayed.

**Figure S2:**
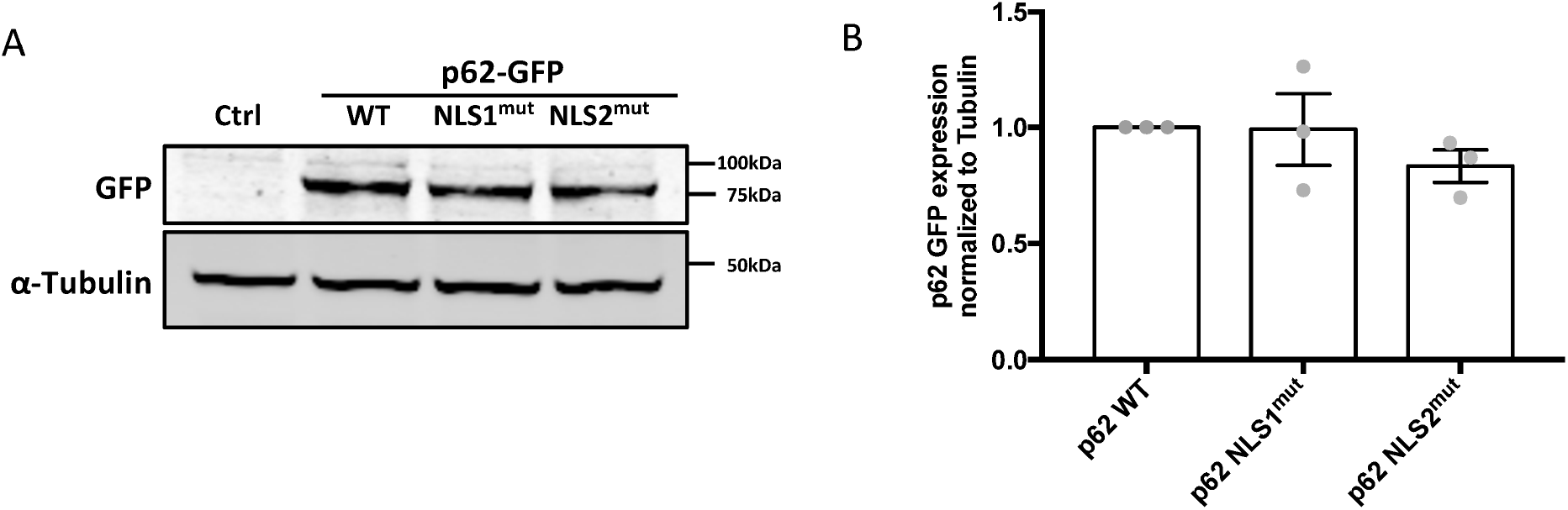
Validation of GFP-p62 overexpression. A) Immunoblot analysis of HeLa cells expressing WT, NLS1^mut^ or NLS2^mut^ p62 at 18 h. Immunoblots directed against tubulin served for normalization. B) Quantification of p62 expression displayed as mean ± SEM. Unpaired T-test with ***P<0.001. Experiment was performed in triplicate and a representative blot displayed.

**Figure S3:**
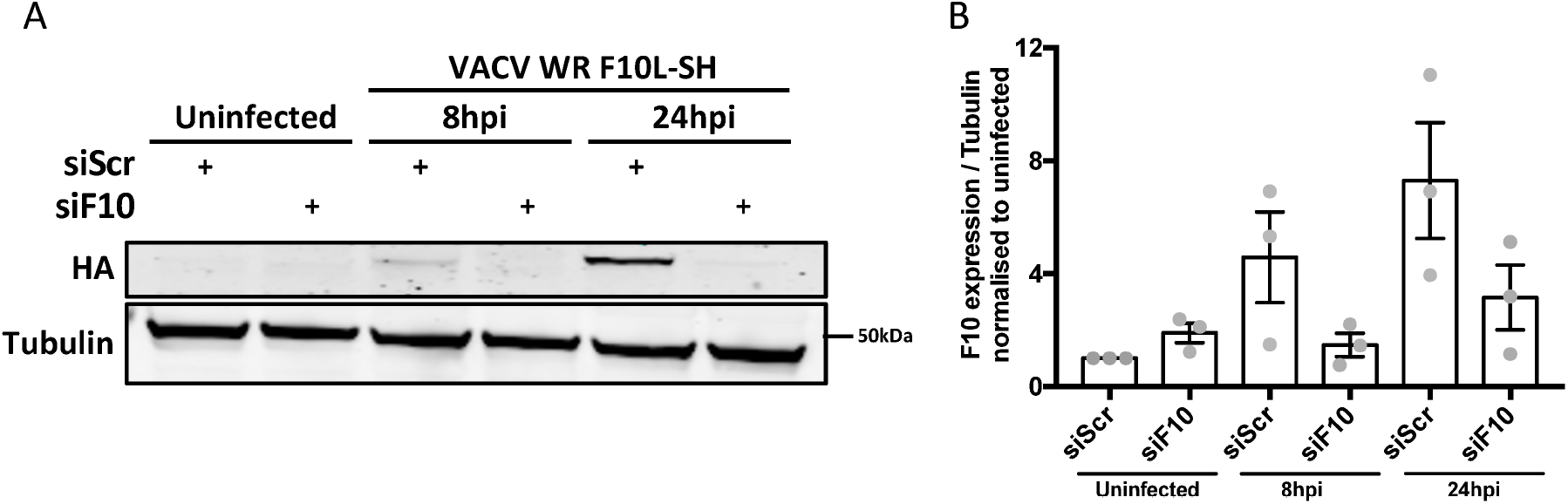
Validation of F10L siRNA. A) HeLa cells transfected with either scrambled or F10-targeting siRNA were infected with VACV F10L-SH. F10 protein levels were determined at 8 and 24 hpi by immunoblot directed against the HA-tag. B) Quantification of F10 protein levels normalized to tubulin are displayed as mean ± SEM. Experiments were performed in triplicate and a representative blot shown.

## Notes

### Competing Interest Statement

The authors have declared no competing interest.

